# G-Protein signaling accelerates stem cell divisions in *Drosophila* males

**DOI:** 10.1101/433623

**Authors:** Manashree Malpe, Leon F. McSwain, Karl Kudyba, Chun L. Ng, Jennie Nicholson, Maximilian Brady, Yue Qian, Vinay Choksi, Alicia G. Hudson, Benjamin B. Parrott, Cordula Schulz

## Abstract

Adult stem cells divide to renew the stem cell pool and replenish specialized cells that are lost due to death or usage. However, little is known about the mechanisms regulating how stem cells adjust to a demand for specialized cells. A failure of the stem cells to respond to this demand can have serious consequences, such as tissue loss, or prolonged recovery post injury.

Here, we challenge the male germline stem cells (GSCs) of *Drosophila melanogaster* for the production of specialized cells using mating experiments. We show that repeated mating reduced the sperm pool and accelerated germline stem cell (GSC) divisions. The increase in GSC divisions depended on the activity of the highly conserved G-proteins. Germline expression of RNA-Interference (RNA-*i*) constructs against G-proteins or a dominant negative G-protein eliminated the increase in GSC divisions in mated males. Consistent with a role for the G-proteins in the regulation of GSC divisions, RNA-*i* against seven out of 35 G-protein coupled receptors (GPCRs) within the germline cells also eliminated the capability of males to accelerate their GSC divisions in response to mating. Our data show that GSCs are receptive to GPCR stimulus, potentially through a network of interactions among multiple signaling pathways.

## Introduction

Metazoan tissues undergo homeostasis wherein stem cells divide and their daughter cells proliferate and differentiate to replace lost cells. The human hematopoietic stem cells, for example, renew a remarkable number of about one trillion blood cells per day ^1, 2^. Stem cells have to maintain a baseline mitotic activity for the production of daughter cells that account for the daily turnover of differentiated cells. However, whether stem cells can modulate their mitotic activity in response to demands that challenge the system is not fully explored. In some instances, stem cells respond to physiological cues; for example, murine hematopoietic stem cells divide more frequently during pregnancy due to increased oestrogen levels ^3^. In *Drosophila melanogaster*, intestinal stem cells initiate extra cell divisions upon ablation of differentiated gut cells. *Drosophila* GSCs modulate their mitotic activity in response to environmental conditions, such as nutrient availability and temperature ^4–7^.

*Drosophila* is an excellent model for identifying the molecules and mechanisms that regulate and fine-tune tissue homeostasis. A plethora of genetic tools are available for manipulating and monitoring dividing adult stem cells in *Drosophila*. The small size of the fly, the short generation cycle, and the fairly low costs covering their maintenance allow for high throughput screens. Here, we subjected several thousand male and several million virgin female flies to mating experiments, a task challenging to perform with vertebrate model organisms. We discovered that repeated mating caused a reproducible and significant increase in GSC division frequency in *Drosophila wild-type* (*wt*) males. Our analysis revealed that this response to mating was dependent on the activity G-proteins. Impairing G-protein activity from the germline cells eliminated the ability of the GSCs to increase their division frequency in response to mating.

G-proteins are highly conserved molecules that associate with GPCRs. GPCRs constitute a large family of cell surface receptors that mediate the cell’s response to a wide range of external stimuli, including odors, pheromones, hormones, and neurotransmitters. Loss of GPCR signaling affects countless developmental and neural processes in humans, as well as vertebrate and invertebrate model organisms ^8–10^. Here we show that reducing the expression of seven out of 35 GPCRs via RNA-*i* from the germline cells eliminated the capability of males to accelerate their GSC divisions when mated. These were the Serotonin (5-HT) Receptors 1A, 1B and 7, Metuselah (Mth), Metuselah-like5 (Mth-l5), Octopamineβ2-Receptor (Octβ2R), and a predicted GPCR encoded by *CG12290*.

A role for any of these GPCRs in regulating GSC division frequency is novel. No previous study has identified any functional role for Mth-l5 or CG12290. Serotonin, Octopamine, and Mth signaling play opposing roles in life-span, locomotion, and sleep ^11–16^. Mth signaling also regulates vesicle trafficking at the synapse, Octopamine signaling regulates ovulation, and Serotonin signaling plays essential roles in memory formation and learning ^17–19^.

## Results

### Mating increased the percentage of GSCs in mitosis

As is typical for many stem cells, the *Drosophila* GSCs are found in a specific cellular microenvironment. They are located at the tip of the gonad, where they are attached to somatic hub cells (Figure 1A, A’). Upon GSC division, one of the daughter cells, called gonialblast, undergoes four rounds of stem cell daughter characteristic transit amplifying divisions, resulting in 16 spermatogonia. Subsequently, spermatogonia enter a tissue-specific differentiation process. They grow in size, undergo the two rounds of meiosis, and develop through extensive morphological changes into elongated spermatids ^20^. According to this tightly controlled homeostasis program, each GSC division can only produce 64 spermatids (Figure 1A). Thus, an increase in sperm production is reliant on the GSCs.

**Figure 1.**
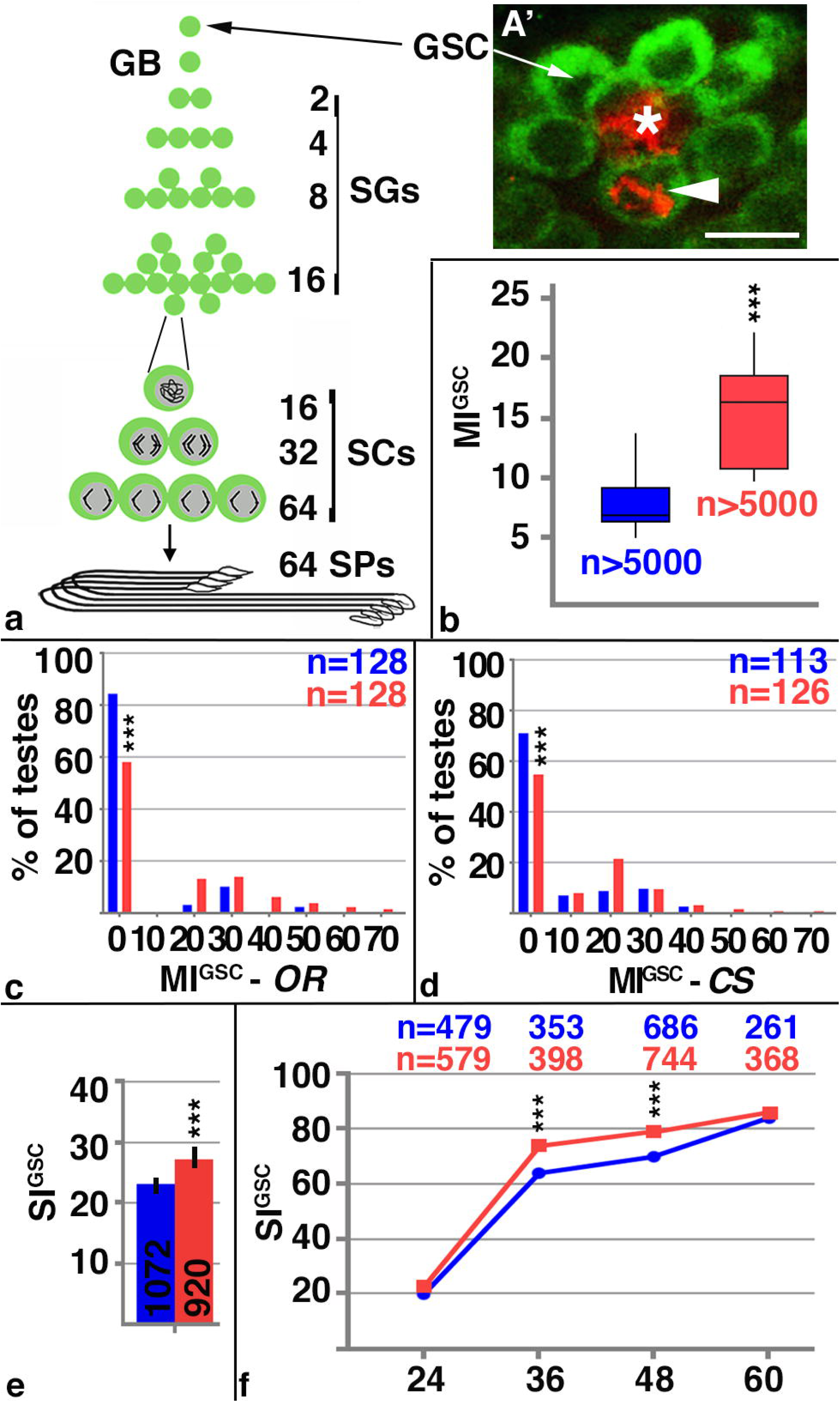
Mating increased male stem cell division frequency. A) Cartoon depicting the stages of *Drosophila* spermatogenesis. Note that every GSC division produces exactly 64 spermatids. GB: gonialblast, SG: spermatogonia, SC: spermatocytes, SP: spermatids. A’) The apical tip of a *wt* testis. The FasIII-positive hub (asterix) is surrounded by seven Vasa-positive GSCs (green), one of which is in mitosis based on anti-pHH3-staining (arrowhead). Scale bar: 10μm. B-F) Blue: non-mated condition, red: mated condition, ***: P-value < 0.001, numbers of GSCs and number of gonads (n=) as indicated. B) Box plots showing the range of MI^GSC^. Lines within boxes represent medians, whiskers represent outliers. C, D) FDGs showing bin of MI^GSC^ (bin width=10) across a population of c) *OR* and D) *CS* males on the X-axis and the percentage of testes with each MI^GSC^ on the Y-axis. E) Bar graph showing SI^GSC^ of *OR* males from three independent experiments. F) Graph showing the percentage of EdU-marked *OR* GSCs on the Y-axis and hours of feeding and mating on the X-axis.

We investigate division frequency using an established immuno-fluorescence protocol ^7^. In this approach, Vasa-positive GSCs are identified based on their position adjacent to FasciclinIII (FasIII)-positive hub cells (Figure 1A’). The percentage of GSCs in mitosis, the M-phase index (MI), is investigated by staining against phosphorylated Histone-H3 (pHH3). The MI of the GSCs (MI^GSC^) is calculated by dividing the number of pHH3-positive GSCs by the total number of GSCs.

To investigate if stem cells can modulate their division frequency in response to a demand for specialized cells, we challenged *Drosophila* males in mating experiments. For each experiment, 80-100 males were exposed individually to virgin females. An equal number of male siblings were each kept in solitude and served as the non-mated controls. To keep experimental variation to a minimum, we employed a three-day mating protocol for all experiments, kept the animals under the same conditions, dissected the testes at the same time of the same day, and dissected experimental groups in tandem. Using *wt* males, we obtained robust and reproducible increases in MI^GSC^ in response to mating. The box-plot in Figure 1B shows the observed difference in MI^GSC^ between mated and non-mated populations of isogenized *wt*, *Oregon R* (*OR*), males from 17 independent mating experiments. Interestingly, we observed variability in MI^GSC^ among males of each condition. The MI^GSC^ of non-mated males ranged from six to nine percent, with a median at seven percent. The MI^GSC^ of mated males ranged from 11 to 18 percent, with a median at 16.5 percent. We hypothesize that this variability in MI^GSC^ within each condition is due to naturally occurring physiological differences within the flies. Likewise, the increase in MI^GSC^ in response to mating varied among the different experiments, but, in each of the experiments, the increase was biologically and statistically significant.

We next investigated if only a few males within a population contributed to the increase in MI^GSC^ or whether the effect of mating is reflected by changes in the MI^GSC^ across a population. These data are displayed in frequency distribution graphs (FDGs). FDGs show how often a particular value is represented within a population. When the distribution of the MI^GSC^ for testes within one population of *OR* flies was plotted, the resulting FDG revealed that mated males had significantly fewer testes with an MI^GSC^ of zero and more testes with higher MI^GSC^ compared to non-mated siblings (Figure 1C). We observed the same result for another isogenized *wt* strain, *Canton S* (*CS*, Figure 1D). We conclude that mating affected the MI^GSC^ of many males within one mated population.

Finally, we asked how long or frequently we had to mate the males to see an increase in MI^GSC^. For this, we mated *OR* males to varying numbers of virgin females and subsequently analyzed how many of their GSCs were in mitotic division. When we exposed *OR* males for 24 hours to one (1F, 24 hrs), two (2F, 24 hrs), or three (3F, 24hrs) female virgins, no significant difference in MI^GSC^ between non-mated and mated males was apparent (Figure S1A). Robust and reproducible increases were seen in *OR* males that were exposed to three virgin females on each of two (2×3F, 48 hrs) or three (3×3F, 72 hrs) days of mating (Figure S1A). We conclude that males have to mate repeatedly for an increase in MI^GSC^ to occur. The increase in MI^GSC^ in mated males was reversible, showing that the response to mating was dynamic. Moving males back into solitude after the three-day mating experiment for 48 hours (3×3F, 120 hrs) eliminated the increase in MI^GSC^ (Figure S1A). Control males mated for 120 hours (5×3F, 120 hrs), in contrast, still had a significant increase in MI^GSC^ (Figure S1A).

### Mating increased GSC division frequency

As another measure of cell divisions, we investigated the percentage of GSCs in synthesis phase (S-phase) of the cell cycle. Testes were labeled with 5-ethynyl-2’-deoxyuridine (EdU) and the S-phase index of the GSCs (SI^GSC^) was calculated by dividing the number of EdU-positive GSCs by the total number of GSCs. Using pulse-labeling experiments, we observed that mated *OR* males displayed significant higher SI^GSC^ compared to their non-mated siblings (Figure 1E). Together with the increase in the MI^GSC^ this suggests that mating accelerates stem cell divisions.

To test this hypothesis, the lengths of the cell cycle were measured using EdU feeding experiments. In this approach, *OR* animals were fed EdU during the mating experiment. We then calculated how many GSCs had been in S-phase at different time points. Our EdU-incorporation experiment revealed that the number of EdU-positive GSCs increased rapidly after 24 hours of feeding and reached 80% at 60 hours of feeding (Figure 1F). Prolonged feeding further increased the numbers of EdU-positive GSCs but this data was excluded from the study as the majority of males that were fed EdU while mating had died by 72 hours of the experiment. The response curve we obtained in this time-course experiment is different from the response curves reported by other groups that used bromo-deoxy-uridine (BrDU) as the thymidine analog instead of EdU. For example, the non-mated males in our experiment had about 70% of EdU-positive GSCs after 48 hours of feeding. A study using *white* (*w*) mutant animals fed the same concentration of the thymidine homologue had a steeper response curve, in which 85% of the GSCs were BrDU-marked after 48 hours of feeding ^21^. Another study using *yellow, vermillion (y, v)* flies showed even steeper response curves where 100% of the GSCs were BrDU-labeled after 24 hours. However, in this study, animals were fed a 30 times higher concentration of the thymidine homologue than used in our study ^22^. We propose that the different response curves are due to the different genetic backgrounds, chemicals and doses.

Most importantly, mated males had significantly more EdU-positive GSCs at 36 and 48 hours of mating compared to their non-mated siblings (Figure 1F). This experiment shows that, in mated males, more GSCs had entered S-phase of the cell cycle. We conclude that mated males had accelerated GSC divisions.

To further investigate how mating affects the cell cycle, we employed the Fly-Fucci technology in combination with the UAS-Ga4 expression system (Duffy, 2002 #321)(Phelps, 1998 #34)(Zielke, 2014 #1122). With Fly-Fucci, the coding regions of fluorescent proteins are fused to the destruction boxes of cell cycle regulators, allowing the marking of different cell cycle stages. These artificial proteins are expressed under control of the Yeast Upstream Activating Sequences ^23^ (Zielke, 2014 #1122). UAS-controlled target genes can be expressed under spatial control using tissue-specific Gal4-transactivators. In addition, temporal control can be applied to their expression by exposing the flies to different temperatures (Phelps, 1998 #34)(Duffy, 2002 #321). For our experiments, we used a *nanos-Gal4*-transactivator (NG4) with a reported expression in GSCs, gonialblasts, and spermatogonia (Van Doren, 1998 #55). Using two independent Fucci-lines, we observed that MI^GSC^ did not significantly increase in mated *Fucci/NG4* males while mated controls animals (*Fucci/wt*) increased their MI^GSC^ compared to non-mated siblings (Figure S1B). We conclude that expressing FUCCI-constructs from these fly lines within the GSCs interfered with their ability to significantly increase MI^GSC^ in response to mating. One possible explanation for this could be that the expression of proteins with destruction boxes could overload the cell cycle machinery of male GSCs.

### Mating reduced the sperm pool

To confirm that our mating experiments created a demand for sperm, we explored differences in the sperm pool of the seminal vesicles between non-mated and mated males. For this, we used two different transgenic constructs that label the sperm. A Don Juan-Green Fluorescent Protein (DJ-GFP) reporter labels the sperm bodies and allows to assess the overall amount of sperm within the seminal vesicles ^24^. A ProtamineB-GFP (Mst35B-GFP) line, on the other hand, only labels the sperm heads and can be used to count the sperm within the seminal vesicles ^25^. With each of these reporters, individualized mature sperm was normally seen within the seminal vesicle of the male reproductive tract.

According to the literature, the total number of sperm within one seminal vesicle varies among different *Drosophila* species and among genetic backgrounds ^25–27^. To keep the genetic background consistent among our experiments, we crossed each of the reporter lines to *OR* females and used their male progeny for our mating experiments. The seminal vesicles were then analyzed at days one to three of the experiment. Based on the size and the fluorescence of the seminal vesicles, we first sorted them into three classes. Class 1 and class 2 seminal vesicles were completely filled with GFP-positive sperm heads. However, class 1 seminal vesicles were very wide (Figure 2A), while class 2 seminal vesicles were thinner (Figure 2B). Class 3 seminal vesicles contained only few GFP-positive sperm heads and had areas that were not filled with GFP (Figure 2C, arrows). A quantification revealed that non-mated males had mostly class 1 and 2 seminal vesicles, while mated males had mostly class 3 seminal vesicles. While we still detected class 1 and 2 seminal vesicles in males that had mated for only one day, their numbers were severely reduced in males after two and three days of mating (Figure 2D-F).

**Figure 2.**
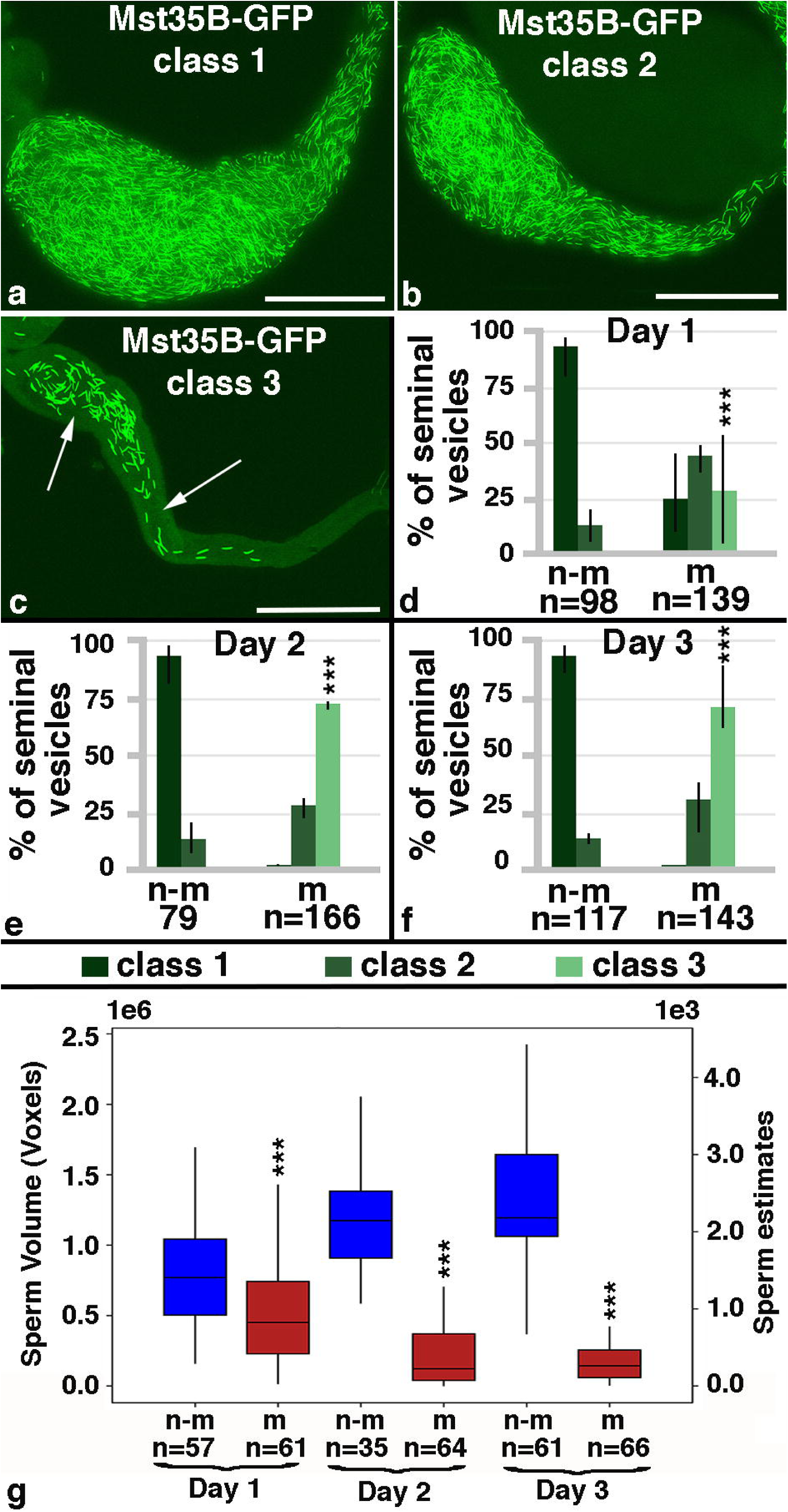
Mating reduced the mature sperm pool. A-C) Class 1,2 and 3 seminal vesicles from Mst35B-GFP males. Scale bars: 0.1 mm; arrows point to GFP-negative regions. D-G) Numbers of seminal vesicles (n=) as indicated, n-m: non-mated, m: mated, ***: P-value < 0.001. D-F) Bar graphs showing the distribution of Class 1 to 3 seminal vesicles in non-mated and mated males at days one to three of the mating experiment. Three fly lines that carry GFP-marked sperm were used: one that carries Dj-GFP (BL#5417), one that carries the Mst35B-GFP (BL#58408), and one line that carries both constructs (BL#58406). G) Box plot showing sperm head volume (based on MST35B-GFP) per seminal vesicle in non-mated and mated males on days one to three of the experiment.

To further validate our observation that mating reduces the amount of sperm, we developed an automated procedure that calculates the volume occupied by Mst35B-GFP-positive sperm heads per seminal vesicle in all focal planes (voxels in Figure 2G). This allowed us to investigate larger numbers of seminal vesicles compared to a previously reported method, in which images through the seminal vesicles were flattened and sperm heads counted by eye ^25^. Furthermore, a computer-based calculation eliminates subjective bias introduced by the investigator. Based on our computer calculation, the sperm heads of mated males occupied significantly less volume within the seminal vesicles than the sperm in non-mated males (Figure 2G). Notably, the total volume occupied by sperm became more reduced with every day of mating. By days two and three of mating it ranged from 0.1 to 0.4 x 10^6^ voxels per seminal vesicle. The non-mated sibling controls, in contrast, maintained a large GFP-occupied volume in their seminal vesicles, with an average of 1.2 x 10^6^ voxels per seminal vesicle. The computer program estimated the numbers of sperm per seminal vesicle of non-mated males around 2000, while males that were mated for two or three days had less than 500 sperm in their seminal vesicles. As our mated males showed a drastic reduction in sperm, we argue that we have created a demand for sperm.

### Mating had no effect on GSC numbers

It was previously reported that females significantly increased the numbers of their GSCs upon mating ^28^. According to the literature, an adult male gonad contains up to twelve GSCs per testis, but the exact number of GSCs per testes appears to vary among different strains and laboratories. One study using a *wt* strain of males reported six to ten GSCs per testis, while other studies using transgenic males in a *w* mutant genetic background reported 8.94 and 12.3 GSCs per testis, respectively ^29–31^. Among our fly lines, we found variation in GSC numbers as well. The distribution of GSCs ranged from one to 14 per testis, with an average of seven GSCs per testis. Males from an isogenized *OR* stock had the lowest average number of GSCs, having only four to five GSCs per testis (Figure 3A). Males from an isogenized *CS* stock had an average of six GSCs per testis (Figure 3B). Animals mutant for *w* alleles, *w^11^*^18^ and *w^1^*, had on average eight and seven GSCs per testis, respectively (Figures 3C, 3D). Males from a *v^1^, y^1^* stock, which serves as the genetic background for many RNA-*i* lines, had the highest average number of GSCs, at 11 GSCs per testis (Figure 3E). We believe that the obtained GSC numbers are specific to the fly lines in our laboratory and do not necessarily reflect the numbers of GSCs in fly stocks of other laboratories.

**Figure 3.**
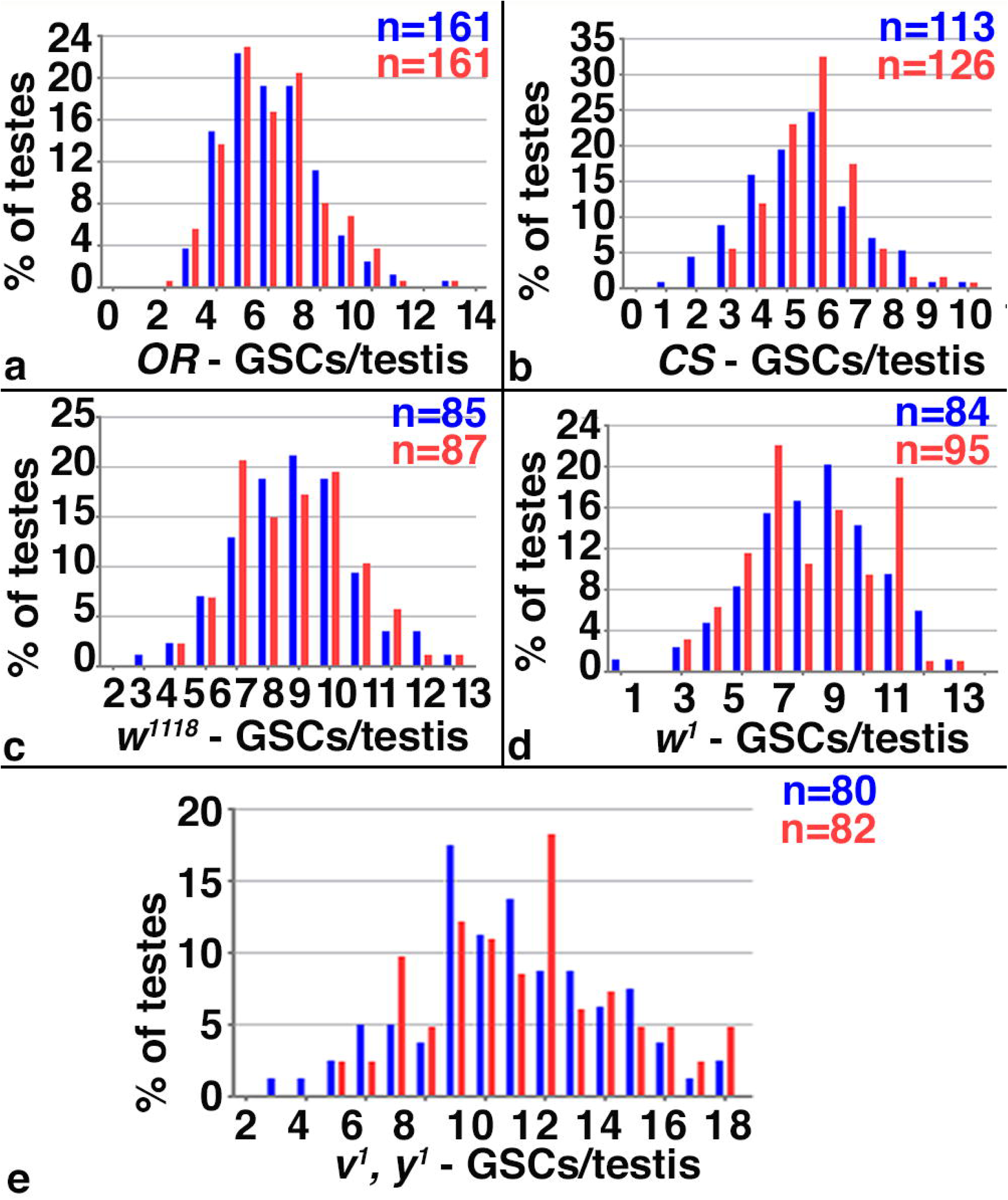
Mating did not affect GSC numbers. A-E) Blue: non-mated condition, red: mated condition, numbers of gonads (n=) as indicated, genotypes as indicated. A-E) FDGs showing numbers of GSCs on the X-axis and percentage of testes with the number of GSCs on the Y-axis. No difference in GSC numbers was observed between non-mated and mated males from different genetic backgrounds.

Importantly, we did not observe a significant difference in the numbers of GSCs between non-mated and mated siblings in any of these fly lines (Figure 3A-E). We concluded that mating did not affect the numbers of GSCs in our fly stocks. However, the observed variation in GSC numbers prompted us to perform our experiments in animals from as similar genetic backgrounds as possible. All males reported in the following of this manuscript carried the X-chromosome from our isogenized *OR* line.

### The increase in MI^GSC^ upon mating required G-protein signaling

*Drosophila* mating is a complex and genetically controlled behavior that is dependent on neural circuits ^32^. This implicates a possible neuronal control in regulating GSC divisions during mating. Therefore, we wanted to focus on the type of signaling pathway commonly stimulated during neural activity, G-protein signaling ^33, 34^. In a non-stimulated cell, a trimeric complex of G-proteins, G_α_, G_β_, and G_γ_ is associated with classical GPCRs (Figure 4A, step 1). When ligand binds to the GPCR, a guanidyl exchange factor within the GPCR becomes activated that exchanges GDP for GTP in the G_α_ subunit. The exchange leads to the dissociation of G_α_ and the G_β/γ_ complex from each other and from the GPCR. Remaining attached to the membrane, G_α_ and G_β/γ_ diffuse along it and activate downstream signal transducers (Figure 4A, step 2) ^35, 36^. Most organisms have multiple genes that encode for each of the G-protein subunits. *Drosophila* has six G_α_, three G_β_, and two G_γ_ proteins, yet only a few examples are available in the literature associating a specific *Drosophila* G-protein with an upstream GPCR^35, 37^.

**Figure 4.**
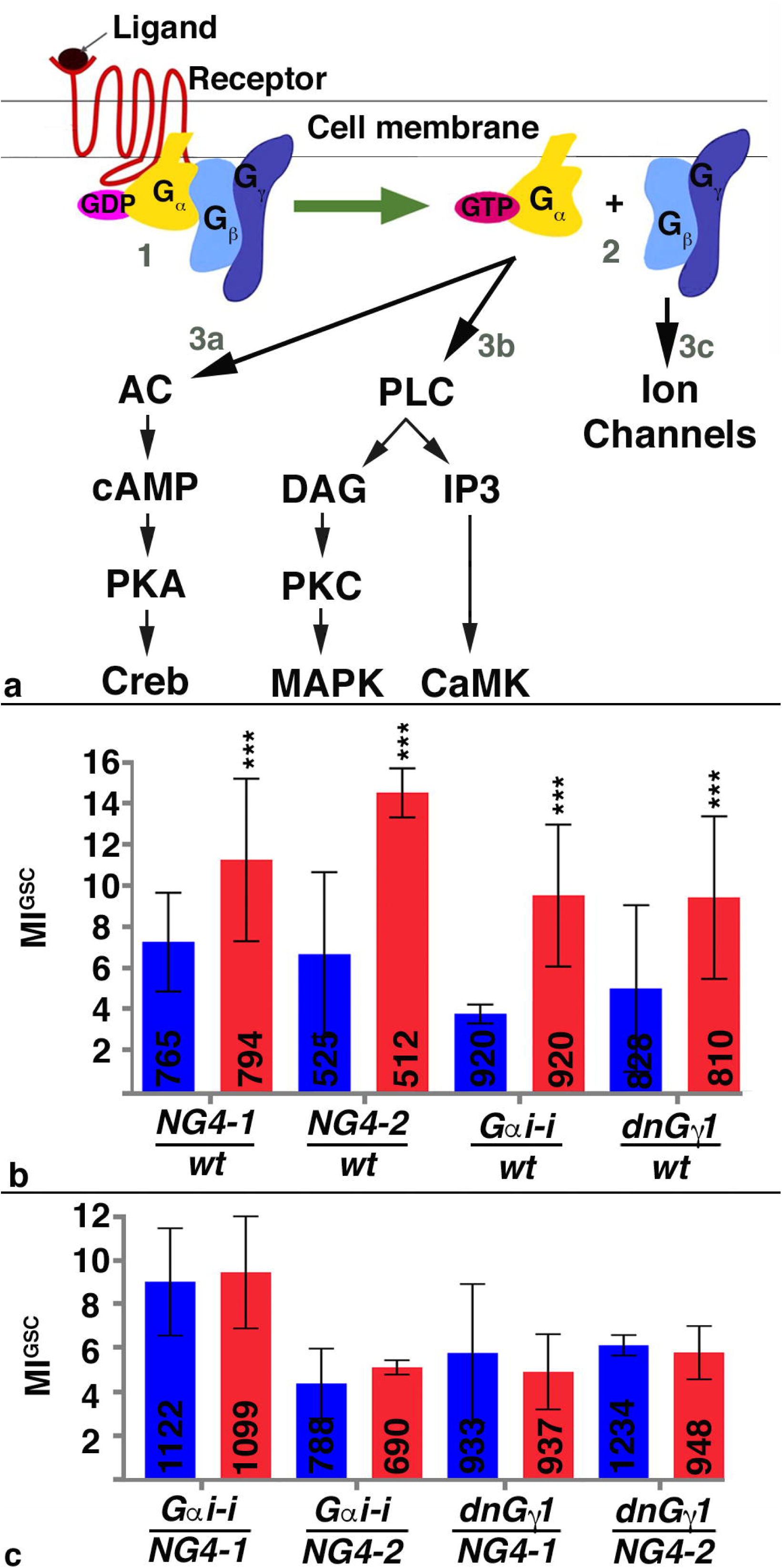
G-proteins were required for the increase in MI^GSC^ in response to mating. A) Cartoon depicting the activation of G-proteins upon GPCR stimulation by ligand. 1: G-protein association before GPCR stimulation, 2: G-protein distribution after GPCR stimulation, 3a-c: downstream signaling cascades. AC: Adenylyl Cyclase, cAMP: cyclic Adenosine Monophosphate, PKA: Protein Kinase A, CREB: cAMP responsive element-binding protein, PLC: Phospho Lipase C, DAG: Diacylglycerol, PKC: Protein Kinase C, MAPK: Map Kinase, IP3: Inositol Triphosphate, CaMK: Calcium^2+/^calmodulin-dependent protein kinase. B, C) Bar graphs showing MI^GSC^. Blue: non-mated condition, red: mated condition, ***: P-value < 0.001, numbers of GSCs as indicated, genotypes as indicated. B) Control animals increased MI^GSC^ in response to mating. C) Males expressing *G*_α_*i-i* or dnG_γ_1 in the germline did not increase MI^GSC^after mating.

Animals mutant for G-protein subunits are often lethal, making it problematic to investigate their roles in the adult. Furthermore, studying G-protein signaling in animals lacking their function throughout the whole body may could affect behavior and physiology of the fly, leading to confounding effects on mating and GSC divisions. Fortunately, large collections of RNA-*i*-lines are available that are expressed under control of UAS. To reduce G-protein signaling we employed two separate *nanos*-Gal4-transactivators (*NG4*), *NG4-1* and *NG4-2.* When RNA-*i* against the different G-protein subunit was expressed within the germline cells via *NG4-1*, several of the mated males displayed only a weak increase in MI^GSC^ compared to their non-mated siblings (Table 1). We focused on an RNA-*i*-line that is directed against the subunit *G_α_i* as animals expressing this construct within the germline did not show any increase in MI^GSC^ in response to mating (Table 1). For reproducibility, we conducted each of the following experiments in triplicates. We used progeny from transgenic Gal4 and UAS-flies that had been crossed to *wt* as positive controls. As expected, each population of positive control males displayed a significant increase in MI^GSC^ when mated (Figure 4B). Experimental flies expressing *G_α_i-i* via *NG4-1* or *NG4-2*, however, failed to increase MI^GSC^ (Figure 4C).

**Table 1.** MI^GSC^ from control, RNA-*i* and overexpression lines directed against G-protein subunits and other signal transducers. UAS-driven expression for the listed genes in the germline via *NG4-1*. BL #: Bloomington stock number, Single and Mated: number of pHH3-positive GSCs/total number of GSCs = MI^GSC^, Diff: MI^GSC^ of mated males minus MI^GSC^ of non-mated males. For RNA-*i*-lines marked by asterisks siblings outcrossed to *wt* did not show a strong response to mating either, suggesting leakiness of the lines.

| Genotype | BL# | Crossed to: | MI<sup>GSC</sup> Single | MI<sup>GSC</sup> Mated | Diff. |
| --- | --- | --- | --- | --- | --- |
| *UAS-G<sub>α</sub>f-i* | 43201 | NG4-1 | 19/448=4.2% | 52/452=11.5% | 7.3 |
|  | 25930* | NG4-1 | 54/1031=5.2% | 76/1183=6.1% | 0.9 |
| *UAS-G<sub>α</sub>i-i* | 34924 | NG4-1 | 22/335=6.6% | 49/322=15.2% | 8.6 |
|  | 40890 | NG4-1 | 33/597=5.5% | 59/536=11% | 5.5 |
|  | 31133 | NG4-1 | 5/269=1.9% | 19/285=6.7% | 4.8 |
| *UAS-G<sub>α</sub>o-i* | 34653* | NG4-1 | 23/313=7.3% | 26/295=8.8% | 1.5 |
|  | 28010 | NG4-1 | 18/240=7.5% | 27/231=11.7% | 4.2 |
| *UAS-G<sub>α</sub>q-i* | 36820 | NG4-1 | 24/403=6.0% | 32/268=11.9% | 5.9 |
|  | 33765 | NG4-1 | 33/320=10% | 50/298=16.8% | 6.8 |
|  | 36775 | NG4-1 | 9/153=5.9% | 21/255=8.6% | 2.7 |
|  | 31268 | OR | 17/233=7.3 | 15/169=8.9 | 1.6 |
|  |  | OR | 14/335=4.2 | 12/164=7.3 | 3.1 |
|  |  | OR | 31/568=5.5 | 27/333=8.1 | 2.6 |
|  |  | NG4-1 | 25/556=4.5 | 8/296=2.7 | -2.8 |
|  |  | NG4-1 | 23/332=6.9 | 23/291=7.9 | 1 |
|  |  | NG4-1 | 23/542=4.2 | 19/577=3.3 | -0.9 |
|  |  | NG4-1 | 71/1430=5.0 | 50/1164=4.3 | -0.7 |
|  | 30735 | NG4-1 | 14/318=4.4% | 23/293=7.8% | 3.4 |
| *UAS-G<sub>α</sub>s-i* | 29576 | NG4-1 | 49/605=8.1% | 60/615=9.7% | 1.6 |
|  | 50704 | NG4-1 | 82/1137=7.3% | 121/1527=7.9% | 0.6 |
| *UAS-G<sub>β</sub>5-i* | 28310 | NG4-1 | 10/306=3.3% | 20/292=6.9% | 3.6 |
| *UAS-G<sub>β</sub>13F-i* | 35041 | NG4-1 | 38/752=4.8% | 46/785=5.7% | 0.9 |
|  | 31134 | NG4-1 | 12/198=6% | 31/221=14% | 8.0 |
| *UAS-G<sub>β</sub>76C-i* | 28507 | NG4-1 | 14/219=6.4% | 26/226=11.5% | 5.1 |
| *UAS-G<sub>γ</sub>1-i* | 25934 | NG4-1 | 21/283=7.4% | 45/311=14.4% | 7.0 |
|  | 34372 | NG4-1 | 21/434=4.8% | 46/400=11.5% | 6.7 |
| *UAS-G<sub>γ</sub>30A-i* | 25932 | NG4-1 | 16/319=5.0% | 18/286=6.3% | 1.3 |
|  | 34484 | NG4-1 | 9/320=2.8% | 31/323=9.6% | 6.8 |
| *UAS-CaMKI-i* | 41900 | NG4-1 | 17/337=5.0% | 31/328=9.4% | 4.4 |
|  | 35362 | NG4-1 | 10/222=4.5% | 17/212=8.0% | 3.5 |
|  | 26726 | NG4-1 | 16/301=5.3% | 22/282=7.8% | 2.5 |
| *UAS-CaMKII-i* | 35330 | NG4-1 | 49/784=6.2% | 91/858=10.6% | 4.4 |
|  | 29401 | NG4-1 | 38/332=11.4% | 43/342=12.6% | 1.2 |
| *UAS-CrebA-i* | 42526 | NG4-1 | 13/301=4.3% | 29/298=9.7% | 5.4 |
| *UAS-Gprk1-i* | 35246 | NG4-1 | 13/323=4.0% | 15/207=7.4% | 3.4 |
|  | 28354 | NG4-1 | 13/304=4.3% | 24/289=12% | 7.7 |
| *UAS-Gprk2-i* | 41933 | NG4-1 | 3/218=1.4% | 24/228=10.5% | 9.1 |
|  | 35326 | NG4-1 | 12/268=4.5% | 35/267=13.1% | 8.6 |
| *UAS-IP3K-i* | 35296 | NG4-1 | 10/225=4.4% | 14/152=9.2% | 4.8 |
|  | 31733 | OR | 25/336=7.4% | 31/331=9.4% | 2.0 |
|  | 31733 | NG4-1 | 67/572=11.7% | 57/555=10.3% | -1.4 |
| *UAS-PKC53E-i* | 34716 | NG4-1 | 18/304=5.9% | 22/289=7.6% | 1.7 |
| *UAS-PKC98E-i* | 29311 | NG4-1 | 14/293=4.8% | 15/284=5.3% | 0.5 |
|  | 35275 | OR | 9/266=3.4% | 25/288=8.4% | 5.0 |
|  |  | NG4-1 | 49/657=7.5% | 45/603=7.5% | 0.0 |
|  | 44074 | OR | 30/597=5.1 | 36/346=10.4 | 5.3 |
|  |  | NG4-1 | 31/289=10.7 | 38/318=11.9 | 1.2 |
| *UAS-PLC2-i* | 33719 | NG4-1 | 23/264=8.7% | 37/311=11.9% | 3.2 |
| *UAS-bsk-i* | 53310 | NG4-1 | 21/380=5.5% | 32/346=9.3% | 3.8 |
| *UAS-lra-i* | 31595 | NG4-1 | 6/272=2.2% | 8/190=4.2% | 2.0 |
| *UAS-kay-i* | 27722 | NG4-1 | 11/256=4.3% | 25/259=9.6% | 5.3 |
|  | 31322 | NG4-1 | 22/384=5.7% | 61/334=18.3% | 12.6 |
|  | 31391 | NG4-1 | 24/291=8.2% | 29/218=13.3% | 5.1 |
| *UAS-rl-i* | 36059 | NG4-1 | 21/269=7.8% | 32/297=10.8% | 3.0 |
| *UAS-wt-5-HT1A* | 27630 | NG4-1 | 16/335=4.8% | 30/302=10% | 5.2 |
|  | 27631 | NG4-1 | 10/240=4.2% | 24/271=8.9% | 4.7 |
| *UAS-wt-CrebB17A* | 7220 | NG4-1 | 51/636=8.0% | 98/628=15.6% | 7.6 |
|  | 9232 | NG4-1 | 32/637=5.0% | 84/592=14.2% | 9.2 |
| *UAS-wt-CaMK2R3* | 29662 | NG4-1 | 20/292=6.8% | 28/289=10.0% | 3.2 |
| *UAS- CaMKII.T287A/* | 29663 | NG4-1 | 16/287=5.6% | 25/283=8.8% | 3.2 |
| *UAS-wt-Gαs* | 6489 | NG4-1 | 32/289=11.1% | 39/300=13.0% | 1.9 |
|  | 6489 | NG4-1 | 40/610=6.6% | 54/652=8.3% | 1.7 |
| *UAS-wt-Ira* | 7216 | NG4-1 | 15/342=4.4% | 44/378=11.6% | 7.2 |
| *UAS-wt-Kay* | 7213 | NG4-1 | 24/341=7.4% | 70/350=20.0% | 12.6 |

We next sought to validate the role for G-protein signaling in GSC division frequency by an alternative approach. A dominant negative version of *Drosophila* G_γ_1 (dnG_γ_1) is available that serves as a reliable tool to abolish G-protein signaling ^38^. Males expressing dnG_γ_1 via either *NG4-1* or *NG4-2* did not show an increase in MI^GSC^ in response to mating (Figure 3C). Control dnG_γ_1/*wt* animals, on the other hand, had increased MI^GSC^ upon mating (Figure 3B). These data clearly show that signaling via G-proteins is required for the increase in MI^GSC^. Plotting the results in FDGs confirmed that mated control animals had significantly fewer testes with an MI^GSC^ of zero and more testes with higher MI^GSC^ compared to non-mated males (Figures S3A-D), and that this response to mating was eliminated in experimental males (Figures S3E-H).

In mammalian cells, three major G-protein-dependent signaling cascades have been described (Figure 3A, steps 3a, b, c) ^33, 39^. For *Drosophila*, the literature provides little information on the signaling cascades downstream of GPCRs but it is generally assumed that the mammalian signal transducers are conserved in flies. To further validate that an increase in MI^GSC^ upon mating is regulated by G-protein signaling we expressed RNA-*i* and mis-expression constructs for conserved signal transducers via NG4 and found that males expressing RNA-*i*-lines for one of the *Drosophila Protein Kinase C* (*PKC*) proteins, *PKC98E*, for *Inositol-triphosphate 3-Kinase* (*IP3K),* and for *Ca2+/Calmodulin-dependent protein kinase II (CaMKII)* indeed failed to increase MI^GSC^ in response to mating (Table 1).

### RNA-i against seven distinct GPCRs blocked the increase in MI^GSC^ upon mating

To further confirm that G-protein signaling regulates the increase in MI^GSC^ we aimed towards identifying the upstream GPCRs. Next Generation Sequencing (NGS) of RNA from *wt* testis tips revealed the expression of 140 receptors, including 35 classical GPCRs (Figure 5 and Table 2). The functions of many of these GPCRs have not been studied yet and mutant animals are only available in rare cases. Expressing RNA-*i*-constructs against most GPCRs in the germline had little to no effect on the ability of the GSCs to increase their MI^GSC^ in response to mating (Table 2). RNA-*i* against three Serotonin Receptors (5HT-1A, 5HT-1B and 5HT-7), Mth, Mth-l5, Octβ2R, and a predicted GPCR encoded by CG12290, clearly and reproducibly eliminated this ability. Animals carrying UAS-controlled RNA-*i*-constructs against these GPCRs (*GPCR-i*) were crossed to *wt*, *NG4-1* and *NG4-2*, and MI^GSC^ of their progeny was investigated. Each of the controls (*GPCR-i/wt*) increased their MI^GSC^ when repeatedly mated to females in each of the triplicate experiments (Figure 5A and Figures S3A-G). However, when the *GPCR-i*-animals were crossed with either *NG4-1* (Figure 5B and Figures S3H-N) or *NG4-2* (Figure 5C and Figures S3O-U) the MI^GSC^ of their non-mated and mated progeny did not significantly differ. Confirming the necessity of the GPCRs in increasing MI^GSC^, we investigated alternative RNA-*i*-lines. A second RNA-*i*-line for Mth blocked the increase in MI^GSC^ in mated males and a second RNA-*i*-line for 5HT-1A displayed only a weak response to mating (Table 3).

**Figure 5.**
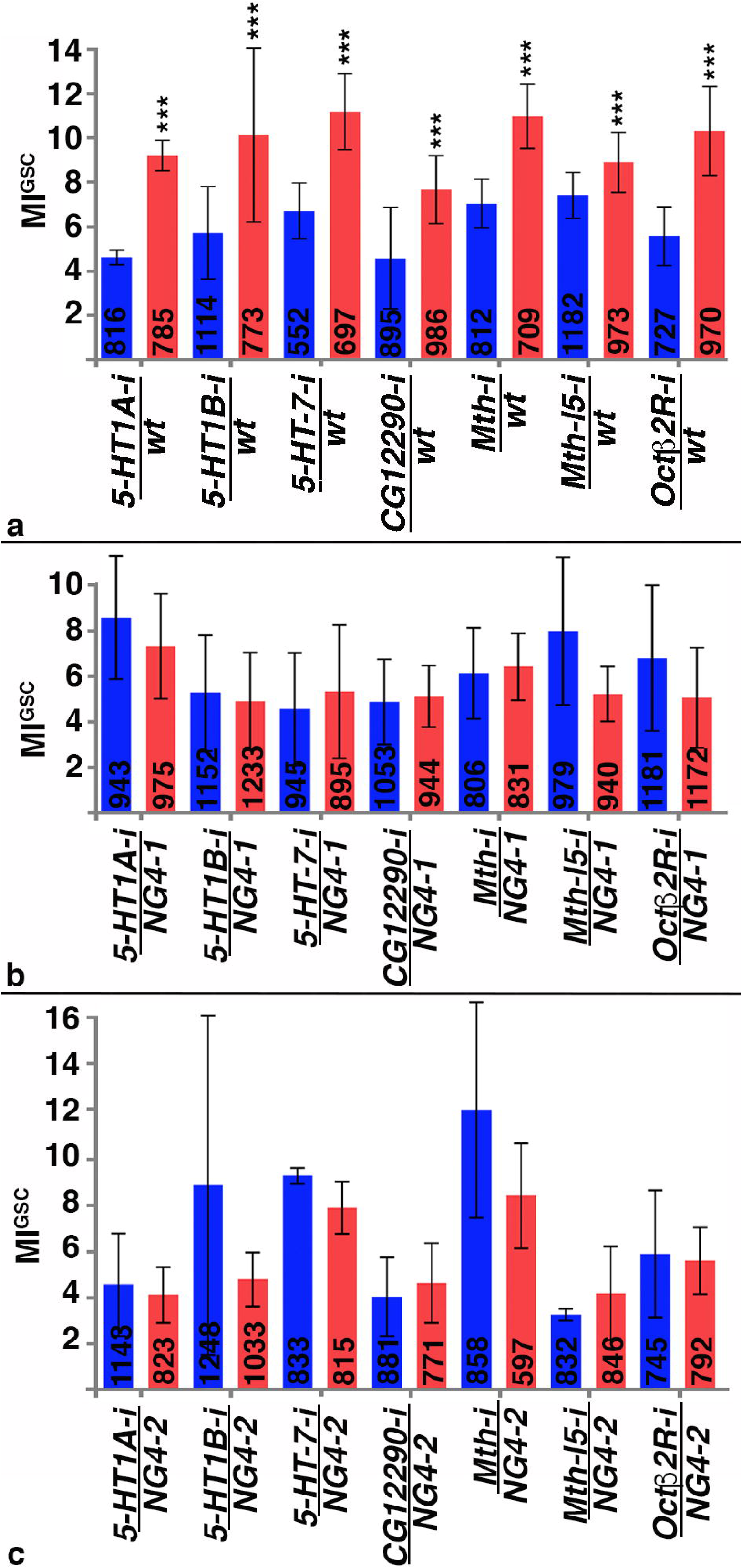
Expression of RNA-*i* against seven distinct GPCRs blocked the increase in MI^GSC^ in response to mating. A-C) Bar graphs showing MI^GSC^. Blue: non-mated condition, red: mated condition, ***: P-value < 0.001, numbers of GSCs as indicated, genotypes as indicated. A) Control males have significantly higher MI^GSC^ than their non-mated siblings. B, C) Mated (B) *GPCR-i*/NG4-1 and (C) *GPCR-i*/NG4-2 males did not have significantly higher MI^GSC^ compared to their non-mated siblings.

**Table 2.** MI^GSC^ from select RNA-*i*-lines directed against GPCRs. UAS-driven expression of RNA-*i* for the listed GPCRs i via *NG4-1* did not block the increase in MI^GSC^ in response to mating. BL #: Bloomington stock number, Single and Mated: number of pHH3-positive GSCs/total number of GSCs = MI^GSC^, Diff: MI^GSC^ of mated males minus MI^GSC^ of non-mated males. Note the variability in MI^GSC^ among the different genotypes. GPCRs marked by asterisks were excluded from further studies because their siblings outcrossed to *wt* did not show a stronger response to mating than the experimental (GPCR-*i*/*NG4-1*) flies.

| GPCR | BL # | Single | Mated | Diff. |
| --- | --- | --- | --- | --- |
| *UAS-5-HT2A-i* | 31882 | 25/490=5.1 | 36/465=7.3 | 2.2 |
|  | 56870 | 19/582=3.3 | 32/553=5.8 | 2.5 |
| *UAS-5-HT2B-i* | 60488 | 6/261=2.3 | 21/272=7.7 | 5.4 |
|  | 25874 | 4/228=1.7 | 34/236=14.4 | 12.7 |
| *UAS-Ado-R-i* | 27536 | 11/276=4.0 | 20/209=9.6 | 5.6 |
| *UAS-AKHR-i* | 29577 | 23/492=4.7 | 54/574=9.4 | 4.7 |
| *UAS-AR-2-i* | 25935 | 13/363=3.6 | 25/336=7.4 | 3.6 |
| *UAS-CG13229-i* | 29519 | 9/285=3.2 | 33/297= 11.1 | 7.9 |
| *UAS-CG14539-i* | 25855 | 21/318=6.6 | 31/307=10.1 | 3.5 |
| *UAS-CG15556-i* | 44574 | 28/425=6.6 | 43/401=10.7 | 4.1 |
| *UAS-CG15744-i* | 28516 | 18/279=6.4 | 27/252=10.7 | 4.3 |
|  | 42497 | 23/323=7.1 | 36/236=11.0 | 3.9 |
| *UAS-CG30106-i* | 27669 | 10/244=4.1 | 34/273=12.4 | 8.3 |
| *UAS-CG33639-i* | 28614 | 32/300=10.7 | 47/371=12.7 | 2.0 |
| *UAS-CCHaR1-i\** | 51168 | 27/407=6.6 | 27/323=8.4 | 1.8 |
| *UAS-Cry-i* | 43217 | 43/389=11.0 | 75/521=14.4 | 3.4 |
| *UAS-CrzR-i* | 52751 | 14/337=4.2 | 31/333=9.3 | 5.1 |
| *UAS-Dop1R1-i* | 62193 | 12/352=3.4 | 25/308=8.1 | 4.6 |
|  | 55239 | 11/300=3.7 | 44/267=16.5 | 12.8 |
| *UAS-GABA*<br>*BR2-i* | 50608 | 6/176=3.4 | 35/207=16.9 | 13.5 |
|  | 27699 | 7/291=2.4 | 18/282=6.4 | 4.0 |
| *UAS-GABA*<br>*BR3-i* | 42725 | 10/190=5.3 | 28/243=11.5 | 6.2 |
| *UAS-Moody-i* | 36821 | 5/301=1.7 | 16/234=6.8 | 5.1 |
| *UAS-Mth-l1-i* | 41930 | 11/279=4.0 | 42/300=14.0 | 10 |
| *UAS-Mth-l3-i* | 41877 | 54/817=6.6 | 81/850=11.8 | 5.2 |
|  | 36822 | 19/231=8.2 | 39/241=16.2 | 8.0 |
| *UAS-Mth-l8-i* | 36886 | 48/933=5.1 | 70/915 =7.6 | 2.5 |
| *UAS-Mth-l9-i* | 51695 | 61/985=6.2 | 82/910 =9.0 | 2.8 |
| *UAS-Mth-l15-i* | 28017 | 14/349=4.0 | 25/337=7.1 | 3.1 |
| *UAS-PK1R-i* | 27539 | 28/478=5.9 | 37/375=9.9 | 4.0 |
| *UAS-Smo-i* | 27037 | 12/263=4.6 | 33/288=11.4 | 6.8 |
|  | 43134 | 19/274=7.0 | 15/147=10.2 | 3.2 |
| *UAS-Tre1-i* | 34956 | 5/234=2.1 | 28/252=11.1 | 9.0 |
| *UAS-TKR86D-i\** | 31884 | 31/564=5.5 | 28/394=7.1 | 1.6 |
| *UAS-TKR99D-i* | 55732 | 30/506=5.9 | 49/467=10.5 | 4.6 |
|  | 27513 | 4/240=1.7 | 30/294=10.2 | 8.5 |

**Table 3.** MI^GSC^ from additional RNA*i*-lines with modified expression of the GPCRs blocking the increase in MI^GSC^ in mated males. BL #: Bloomington stock number, Single and Mated: number of pHH3-positive GSCs/total number of GSCs = MI^GSC^, Diff: MI^GSC^ of mated males minus MI^GSC^ of non-mated males.

| GPCR | BL # | Single | Mated | Diff. |
| --- | --- | --- | --- | --- |
| *5HT-1A-i/NG4-1* | 25834 | 64/841=7.6 | 67/777=8.6 | 1 |
| *5HT-1B-i/NG4-1* | 25833 | 16/256=6.2 | 26/268=9.7 | 3.5 |
|  | 27635 | 3/304=1.0 | 27/317=8.5 | 7.5 |
|  | 51842 | 20/381=5.2 | 32/375=8.5 | 3.3 |
|  | 54006 | 13/405=3.2 | 23/351=6.5 | 3.3 |
| *5HT-7-i/NG4-1* | 32471 | 8/238=3.4 | 17/229=7.4 | 4.0 |
| *CG12290-i/NG4-1* | 42520 | 4/260=1.5 | 31/246=12.6 | 11.1 |
| *Mth-i/NG4-1* | 27495 | 14/352=4 | 15/336=4.5 | 0.5 |

Mating success was evaluated by two criteria: visual observation and the appearance of progeny. When flies were anesthetized to exchange the females for fresh virgins, several copulating pairs of males and females were always observed. Furthermore, 100 single females that had been exposed to males on day one of the experiment were placed into one food vial each and mating success evaluated a few days later by counting the percentage of vials with progeny. Most males in this study sired 60-90% of the females. Specifically, each of the *GPCR-i/NG4-1* males produced offspring (Table S1), showing that a block in the increase in MI^GSC^ is not caused by a failure to mate but by a lack of GPCR signaling. Viable alleles of 5HT-1A and 5HT-1B were not pursued as alternative strategies because they displayed only a weak mating success rate (Table S1).

Finally, we wanted to assure that male age had no effect on the increase in MI^GSC^. We performed a time-course experiment of one, two, three, and four-week old *OR* males. We found that mated males of all ages showed robust increases in MI^GSC^ compared to their non-mated siblings (Figure S1C). We conclude that aging animals for up to four weeks had little to no effect on the ability of *wt* GSCs to increase their MI^GSC^ in response to mating, and that the age of the transgenic animals used in this study (three weeks of age at the time of testes dissection) had no impact on the obtained results.

## Discussion

Here, we show that repeated mating reduced the sperm pool and increased GSC division frequency. Using highly controlled experiments, we demonstrate that mated males had more GSCs in M-phase and S-phase of the cell cycle compared to non-mated males. Mated males also showed faster incorporation of EdU indicating that their GSCs progressed faster through the cell cycle. Our findings demonstrate that GSCs can respond to a demand for sperm by accelerating their mitotic activity. Based on RNA-*i* targeting G-proteins and a dominant negative construct against G_γ_1, the increase in MI^GSC^ of mated males is dependent on G-protein signaling. Furthermore, signal transducers predicted to act downstream of G-proteins and GPCRs predicted to act upstream of G-proteins also appeared to be required for the response to mating.

Due to the lack of mutants and a potential interference of whole animal knock-down in the behavior of the flies, we used tissue-specific expression of RNA-*i*-constructs. It is surprising that our studies revealed potential roles for seven instead of a single GPCR in the increase of MI^GSC^ in response to mating. A possible explanation is that some of the RNA-*i*-lines have off-target effects. RNA*-i*-hairpins can cause the down-regulation of unintended targets due to stretches of sequence homologies, especially when long hairpins are used ^40, 41^. However, with the exception of the RNA-*i*-line directed against 5HT-7, all lines that produced a phenotype contain second generation vectors with a short, 21 nucleotide hairpin predicted to have no off-target effects ^42^. We hypothesize that multiple GPCRs regulate the increase in MI^GSC^ in response to mating. Consistent with this, expression of second RNA-*i*-line directed against Mth or 5HT-1A interfered with the increase in MI^GSC^ in mated males.

Our finding that RNA-*i* against several GPCRs blocked the increase in MI^GSC^ in mated males suggests a high level of complexity in the regulation of GSC divisions. In the literature, increasing evidence has emerged that GPCRs can form dimers and oligomers and that these physical associations have a variety of functional roles, ranging from GPCR trafficking to modification of G-protein mediated signaling ^43–45^. In *C. elegans,* two Octopamine receptors, SER-3 and SER-6, additively regulate the same signal transducers for food-deprived-mediated signaling. One possible explanation for the non-redundant function of the two receptors was the idea that they form a functional dimer ^46^. In mammalian cells, 5-HT receptors can form homo-dimers and hetero-dimers and, dependent on this, have different effects on G-protein signaling ^47–49^. In cultured fibroblast cells, for example, G-protein coupling is more efficient when both receptors within a 5-HT4 homo-dimer bind to agonist instead of only one ^50^. In cultured hippocampal neurons, hetero-dimerization of 5-HT1A with 5-HT7 reduces G-protein activation and decreases the opening of a potassium channel compared to 5-HT1A homo-dimers ^51^. The formation of hetero-dimers of GPCRs with other types of receptors plays a role in depression and in the response to hallucinogens in rodents ^52, 53^.

Alternatively, or in addition to the possibility that some or all of the seven GPCRs form physical complexes, a role for several distinct GPCRs in regulating GSC division frequency could be explained by crosstalk among the downstream signaling cascades. One signaling cascade could, for example, lead to the expression of a kinase that is activated by another cascade. Similarly, one signaling cascade could open an ion channel necessary for the activity of a protein within another cascade. Unfortunately, the literature provides little information on *Drosophila* GPCR signal transduction cascades and only very few mutants have been identified that affect a process downstream of GPCR stimulation. Thus, it remains to be explored how stimulation of the GPCRs and G-proteins increase GSC divisions.

The role for G-protein signaling in regulating the frequency of stem cell divisions is novel. Our data suggest that the increase in MI^GSC^ in response to mating is regulated by external signals, potentially arising from the nervous system, that stimulate G-protein signaling within the GSCs. Based on the nature of the GPCRs, the activating signal could be Serotonin, the Mth ligand, Stunted, Octopamine, or two other, yet unknown, signals that activate Mth-l5, and CG12290 ^54–56^. It will be interesting to address which of these ligands are sufficient to increase MI^GSC^, in what concentrations they act, by which tissues they are released, and whether they also affect other stem cell populations.

## Methods

### Fly husbandry

Flies were raised on a standard cornmeal/agar diet and maintained in temperature-, light-, and humidity-controlled incubators. Unless otherwise noted, all mutations, markers, and transgenic lines are described in the *Drosophila* database and were obtained from the Bloomington stock center (Consortium, 2003 #132).

### UAS/Gal4-expression studies

Two separate X; UAS-*dicer*; nanos-*Gal4* (NG4-1 and NG4-2) fly lines were used as transactivators. Females from the transactivator line or *wt* females were crossed with males carrying target genes under the control of UAS in egg lay containers with fresh apple juice-agar and yeast paste to generate either experimental or control flies. The progeny were transferred into food bottles, raised to adulthood at 18°C, males collected and then shifted to 29°C for seven days to induce high activity of Gal4 prior to the mating experiment. Note that the males were not collected as virgins as to avoid any potential developmental or learning effects on our experiments.

### Mating experiments

Unless otherwise noted, mating experiments were performed at 29°C. Males and virgin females were placed on separate apple juice-agar plates with yeast paste overnight to assure they were well fed prior to their transfer into mating chambers. Single males were placed into each mating slot either by themselves (non-mated) or with three virgin females (mated) and the chambers closed with apple juice-agar lids with yeast paste. Females were replaced by virgin females on each of the following two days and apple juice-agar lids with yeast paste were replaced on a daily basis for both non-mated and mated animals. In most instances, females from the stock *X□X*, *y, w, f* / Y / *shi ^ts^* were L used as virgins. When raised at 29°C, only females hatch from this stock. For fertility tests, *OR* virgins were used. Note that 10-20% of the mated males died during the experiment while only 5% of the non-mated siblings died.

### Immuno-fluorescence and microscopy

Animals were placed on ice to immobilize them. Gonads were dissected in Tissue Isolation Buffer (TIB) and collected in a 1.5 ml tube with TIB buffer on ice for no more than 30 minutes. Gonads were then fixed, followed by immuno-fluorescence staining and imaging as previously described ^7^. The mouse anti-FasciclinIII (FasIII) antibody (1:10) developed by C. Goodman was obtained from the Developmental Studies Hybridoma Bank, created by the NICHD of the NIH and maintained at The University of Iowa, Department of Biology, Iowa City, IA 52242. Goat anti-Vasa antibody (1:50 to 1:300) was obtained from Santa Cruz Biotechnology Inc. (sc26877), anti-phosphorylated Histone H3 (pHH3) antibodies (1:100 to 1:1000) were obtained from Fisher (PA5-17869), Milllipore (06-570), and Santa Cruz Biotechnology Inc. (sc8656-R). Secondary Alexa 488, 568, and 647-coupled antibodies (1:1000) and Slow Fade Gold embedding medium with DAPI were obtained from Life Technologies. Images were taken with a Zeiss Axiophot, equipped with a digital camera, an apotome, and Axiovision Rel. software. Statistical relevance was determined using the two-tailed Graphpad student’s t-test.

### EdU-labeling experiments

The EdU-labeling kit was obtained from Invitrogen and the procedure performed following manufacturer’s instructions. For EdU-pulse labeling experiments, animals were mated as described above, and the dissected testes incubated with 10mM EdU in PBS for 30 minutes at room temperature prior to fixation. For EdU-feeding experiments, *OR* males were fed 10 mM EdU in liquid yeast provided on paper towels. These animals were mated at room temperature (21°C) because the paper towels easily dried out at higher temperatures, causing the flies to dehydrate and die.

### Sperm head volumetric calculations

In order to easily evaluate sperm numbers, we turned to computer analysis in Python. By quantifying the volume of GFP signal we generated estimates to the amount of sperm in each seminal vesicle. Image stacks were taken of individual seminal vesicles. After masking relevant regions, each image set was normalized by mean subtraction and division by the standard deviation, followed by rescaling image intensity to encompass the range of the image. To remove signal noise, a median filter was applied and the mask refined by Otso thresholding. We determined signal volume by hysteresis thresholding. This approach initially thresholds an image at an upper limit, and then expands the region by adjacent pixels satisfying the lower threshold. We set the lower bound at the value generated from a triangle threshold and the upper threshold as the median value above the lower limit. The number of signal voxels was calculated and normalized to an expected size of a single sperm head. Our analysis utilized OpenCV 3.4.2, Scipy1.2.1, Scikit-image 0.14.2, Numpy 1.16.2, Matplotlib 3.0.3, Seaborn 0.9.0, as well as built in Python 3.7.3 modules ^57–61^.

## Acknowledgements

The authors dedicate this manuscript to Bruce Baker, who was one of the foremost scientists in the field, and a great colleague and friend. The authors are grateful to Richard Zoller, Yue Qian, Megan Aarnio, Heather Kudyba, Jacqueline Uribe, Chidemman Ihenacho, Stefani Moore, Sampreet Reddy, Amanda Cameron, Chederli Belongilot, Dylan Ricke, Kenneth Burgess, Amanda Redding, Erin Guillebeau, Jennifer Murphy, Chantel McCarty, Sarah Murphy, Haley Grable, Mitch Hanson, Haein Kim, and Sarah Rupert for technical assistance. We thank Bruce Baker, Carmen Robinett, Edward Kravitz, Matthew Freeman, Mark Brown, Erika Matunis, Steve DiNardo, Margaret Fuller, Alan Spradling, Hannele Rahuola-Baker, Eric Bohman, Celeste Berg, Wolfgang Lukowitz, Patricia Moore, Jim Lauderdale, Michael Tiemeyr, Scott Dougan, Rachel Roberts-Galbraith, and Rhonda Snook for helpful discussions, and Heath Aston Zachary Letts for comments on the manuscript. We are especially grateful to Barry Ganetzky for the *X□X, shi ^ts^* fly stock and to Wolfgang Lukowitz for the use of his L microscope. This work was supported by NSF grants #0841419 and #1355009, and UGA bridge funds given to CS.

## Author contributions

M.M, B.B.P, and C.S developed and supervised the project, L.F.M coordinated the mating experiments, K.K. identified the GPCRs expressed in testes tips and developed the computer analysis for the sperm counts, all authors performed the experiments, M.M, K.K., and CS wrote the manuscript.

## References

1. Dancey, J.T., Deubelbeiss, K.A., Harker, L.A. & Finch, C.A. Neutrophil kinetics in man. J Clin Invest 58, 705–715 (1976).

2. Erslev, A. Production of erythrocytes, in Hematology. (ed. B.E. William WJ, Erslev AJ, Lichtman MA) 365–376 (Mc-Graw-Hill, New York, NY; 1983).

3. Nakada, D. et al. Oestrogen increases haematopoietic stem-cell self-renewal in females and during pregnancy. Nature 505, 555–558 (2014).

4. Hsu, H.J., LaFever, L. & Drummond-Barbosa, D. Diet controls normal and tumorous germline stem cells via insulin-dependent and -independent mechanisms in Drosophila. Dev Biol 313, 700–712 (2008).

5. Amcheslavsky, A., Jiang, J. & Ip, Y.T. Tissue damage-induced intestinal stem cell division in Drosophila. Cell Stem Cell 4, 49–61 (2009).

6. McLeod, C.J., Wang, L., Wong, C. & Jones, D.L. Stem cell dynamics in response to nutrient availability. Curr Biol 20, 2100–2105 (2010).

7. Parrott, B.B., Hudson, A., Brady, R. & Schulz, C. Control of germline stem cell division frequency--a novel, developmentally regulated role for epidermal growth factor signaling. PLoS One 7, e36460 (2012).

8. Schoneberg, T. et al. Mutant G-protein-coupled receptors as a cause of human diseases. Pharmacol Ther 104, 173–206 (2004).

9. Wettschureck, N. & Offermanns, S. Mammalian G proteins and their cell type specific functions. Physiol Rev 85, 1159–1204 (2005).

10. Langenhan, T. et al. Model Organisms in G Protein-Coupled Receptor Research. Mol Pharmacol 88, 596–603 (2015).

11. Lin, Y.J., Seroude, L. & Benzer, S. Extended life-span and stress resistance in the Drosophila mutant methuselah. Science 282, 943–946 (1998).

12. Selcho, M., Pauls, D., El Jundi, B., Stocker, R.F. & Thum, A.S. The role of octopamine and tyramine in Drosophila larval locomotion. J Comp Neurol 520, 3764–3785 (2012).

13. Silva, B., Goles, N.I., Varas, R. & Campusano, J.M. Serotonin receptors expressed in Drosophila mushroom bodies differentially modulate larval locomotion. PLoS One 9, e89641 (2014).

14. Crocker, A. & Sehgal, A. Octopamine regulates sleep in drosophila through protein kinase A-dependent mechanisms. J Neurosci 28, 9377–9385 (2008).

15. Yuan, Q., Joiner, W.J. & Sehgal, A. A sleep-promoting role for the Drosophila serotonin receptor 1A. Curr Biol 16, 1051–1062 (2006).

16. Li, Y. et al. Octopamine controls starvation resistance, life span and metabolic traits in Drosophila. Sci Rep 6, 35359 (2016).

17. Song, W. et al. Presynaptic regulation of neurotransmission in Drosophila by the g protein-coupled receptor methuselah. Neuron 36, 105–119 (2002).

18. Lee, H.G., Seong, C.S., Kim, Y.C., Davis, R.L. & Han, K.A. Octopamine receptor OAMB is required for ovulation in Drosophila melanogaster. Dev Biol 264, 179–190 (2003).

19. Sitaraman, D. et al. Serotonin is necessary for place memory in Drosophila. Proc Natl Acad Sci U S A 105, 5579–5584 (2008).

20. Fuller, M.T. Spermatogenesis, in The development of Drosophila melanogaster, Vol. 1. (ed. M. Bate, Martinez-Arias, A) 71–147 (Cold Spring Harbor Press, Cold Spring Harbor; 1993).

21. Wallenfang, M.R., Nayak, R. & DiNardo, S. Dynamics of the male germline stem cell population during aging of Drosophila melanogaster. Aging Cell 5, 297–304 (2006).

22. Yang, H. & Yamashita, Y.M. The regulated elimination of transit-amplifying cells preserves tissue homeostasis during protein starvation in Drosophila testis. Development 142, 1756–1766 (2015).

23. Abdouh, M., Albert, P.R., Drobetsky, E., Filep, J.G. & Kouassi, E. 5-HT1A-mediated promotion of mitogen-activated T and B cell survival and proliferation is associated with increased translocation of NF-kappaB to the nucleus. Brain Behav Immun 18, 24–34 (2004).

24. Santel, A., Blumer, N., Kampfer, M. & Renkawitz-Pohl, R. Flagellar mitochondrial association of the male-specific Don Juan protein in Drosophila spermatozoa. J Cell Sci 111 (Pt 22), 3299–3309 (1998).

25. Tirmarche, S. et al. Drosophila protamine-like Mst35Ba and Mst35Bb are required for proper sperm nuclear morphology but are dispensable for male fertility. G3 (Bethesda) 4, 2241–2245 (2014).

26. Pitnick, S., Markow, T.A. Male gametic Strategies: Sperm Size, Testes Size, and the Allocation of Ejaculate among Successive Mates by the Sperm-Limited Fly Drosophila Pachea and its Relatives. The American Naturalist 143, 785–819 (1994).

27. Kubrak, O.I., Kucerova, L., Theopold, U., Nylin, S. & Nassel, D.R. Characterization of Reproductive Dormancy in Male Drosophila melanogaster. Front Physiol 7, 572 (2016).

28. Ameku, T. & Niwa, R. Mating-Induced Increase in Germline Stem Cells via the Neuroendocrine System in Female Drosophila. PLoS Genet 12, e1006123 (2016).

29. Chen, D. et al. Gilgamesh is required for the maintenance of germline stem cells in Drosophila testis. Sci Rep 7, 5737 (2017).

30. Yamashita, Y.M., Jones, D.L. & Fuller, M.T. Orientation of asymmetric stem cell division by the APC tumor suppressor and centrosome. Science 301, 1547–1550 (2003).

31. Sheng, X.R. & Matunis, E. Live imaging of the Drosophila spermatogonial stem cell niche reveals novel mechanisms regulating germline stem cell output. Development 138, 3367–3376 (2011).

32. Manoli, D.S., Fan, P., Fraser, E.J. & Shah, N.M. Neural control of sexually dimorphic behaviors. Curr Opin Neurobiol 23, 330–338 (2013).

33. Geppetti, P., Veldhuis, N.A., Lieu, T. & Bunnett, N.W. G Protein-Coupled Receptors: Dynamic Machines for Signaling Pain and Itch. Neuron 88, 635–649 (2015).

34. Lee, D. Global and local missions of cAMP signaling in neural plasticity, learning, and memory. Front Pharmacol 6, 161 (2015).

35. McCudden, C.R., Hains, M.D., Kimple, R.J., Siderovski, D.P. & Willard, F.S. G-protein signaling: back to the future. Cell Mol Life Sci 62, 551–577 (2005).

36. Oldham, W.M. & Hamm, H.E. Heterotrimeric G protein activation by G-protein-coupled receptors. Nat Rev Mol Cell Biol 9, 60–71 (2008).

37. Boto, T., Gomez-Diaz, C. & Alcorta, E. Expression analysis of the 3 G-protein subunits, Galpha, Gbeta, and Ggamma, in the olfactory receptor organs of adult Drosophila melanogaster. Chem Senses 35, 183–193 (2010).

38. Deshpande, G., Godishala, A. & Schedl, P. Ggamma1, a downstream target for the hmgcr-isoprenoid biosynthetic pathway, is required for releasing the Hedgehog ligand and directing germ cell migration. PLoS Genet 5, e1000333 (2009).

39. Moolenaar, W.H. G-protein-coupled receptors, phosphoinositide hydrolysis, and cell proliferation. Cell Growth Differ 2, 359–364 (1991).

40. Kulkarni, M.M. et al. Evidence of off-target effects associated with long dsRNAs in Drosophila melanogaster cell-based assays. Nat Methods 3, 833–838 (2006).

41. Moffat, J., Reiling, J.H. & Sabatini, D.M. Off-target effects associated with long dsRNAs in Drosophila RNAi screens. Trends Pharmacol Sci 28, 149–151 (2007).

42. Perkins, L.A. et al. The Transgenic RNAi Project at Harvard Medical School: Resources and Validation. Genetics 201, 843–852 (2015).

43. Filizola, M. & Weinstein, H. The study of G-protein coupled receptor oligomerization with computational modeling and bioinformatics. FEBS J 272, 2926–2938 (2005).

44. Milligan, G. G protein-coupled receptor dimerisation: molecular basis and relevance to function. Biochim Biophys Acta 1768, 825–835 (2007).

45. Terrillon, S. & Bouvier, M. Roles of G-protein-coupled receptor dimerization. EMBO Rep 5, 30–34 (2004).

46. Yoshida, M., Oami, E., Wang, M., Ishiura, S. & Suo, S. Nonredundant function of two highly homologous octopamine receptors in food-deprivation-mediated signaling in Caenorhabditis elegans. J Neurosci Res 92, 671–678 (2014).

47. Lukasiewicz, S. et al. Hetero-dimerization of serotonin 5-HT(2A) and dopamine D(2) receptors. Biochim Biophys Acta 1803, 1347–1358 (2010).

48. Herrick-Davis, K. Functional significance of serotonin receptor dimerization. Exp Brain Res 230, 375–386 (2013).

49. Xie, Z., Lee, S.P., O’Dowd, B.F. & George, S.R. Serotonin 5-HT1B and 5-HT1D receptors form homodimers when expressed alone and heterodimers when co-expressed. FEBS Lett 456, 63–67 (1999).

50. Pellissier, L.P. et al. G protein activation by serotonin type 4 receptor dimers: evidence that turning on two protomers is more efficient. J Biol Chem 286, 9985–9997 (2011).

51. Renner, U. et al. Heterodimerization of serotonin receptors 5-HT1A and 5-HT7 differentially regulates receptor signalling and trafficking. J Cell Sci 125, 2486–2499 (2012).

52. Borroto-Escuela, D.O., Tarakanov, A.O. & Fuxe, K. FGFR1-5-HT1A Heteroreceptor Complexes: Implications for Understanding and Treating Major Depression. Trends Neurosci 39, 5–15 (2016).

53. Moreno, J.L., Holloway, T., Albizu, L., Sealfon, S.C. & Gonzalez-Maeso, J. Metabotropic glutamate mGlu2 receptor is necessary for the pharmacological and behavioral effects induced by hallucinogenic 5-HT2A receptor agonists. Neurosci Lett 493, 76–79 (2011).

54. Saudou, F., Boschert, U., Amlaiky, N., Plassat, J.L. & Hen, R. A family of Drosophila serotonin receptors with distinct intracellular signalling properties and expression patterns. EMBO J 11, 7–17 (1992).

55. Cvejic, S., Zhu, Z., Felice, S.J., Berman, Y. & Huang, X.Y. The endogenous ligand Stunted of the GPCR Methuselah extends lifespan in Drosophila. Nat Cell Biol 6, 540–546 (2004).

56. Maqueira, B., Chatwin, H. & Evans, P.D. Identification and characterization of a novel family of Drosophila beta-adrenergic-like octopamine G-protein coupled receptors. J Neurochem 94, 547–560 (2005).

57. van der Walt, S., Colbert, C., Varoquaux, G. The NumPy Array: A Structure for Efficient Numerical Vomputation. Computing in Science and Engineering, 22–30 (2011).

58. van der Walt, S., Schoenberger, J.L., Nunez-Iglesias, J., Boulogne, F., Warner, J.D., Yager, N., Gouillart, E., Yu, T., and the scikit-image contributors scikit-image: Image processing in Python. PerJ:e453 (2014).

59. Travis, E., Oliphant, E. A guide to NumPy. (2006).

60. Hunter, J.D. Matplotib: A 2D Graphics Environment. Computing in Science and Engineering, 90–95 (2007). https://archive.org/details/NumPyBook.

61. Jones, E., Oliphant, E., Peterson, P., et al SciPy: Open Source Scientific Tools for Python. (2001-). https://www.scipy.org.

