## Supplemental table and figures for "G-Protein signaling accelerates stem cell divisions in *Drosophila* males"

**Table S1. Fertility Assay**

Male fertility was calculated based on the % of females that produced offspring after mating with males of the indicated genotype. BL#: Bloomington stock number.

| **Genotype** | **BL #** | **Male fertility** |
| --- | --- | --- |
| *OR* | N/A | 72% |
| *CS* | N/A | 62% |
| *5HT-1A-i /NG4-1* | 33885 | 75% |
| *5HT-1B-i /NG4-1* | 33418 | 61% |
| *5HT-7-i/NG4-2* | 27273 | 91% |
| *CG12290-i /NG4-1* | 31873 | 77% |
| *Mth-i /NG4-1* | 36823 | 90% |
| *Mth-l5-i /NG4-1* | 42515 | 84% |
| *Octβ2R-i /NG4-1* | 50580 | 89% |
| *5HT-1A^Δ5kb^ /5HT-1A^Δ5kb^* | 27640 | 15% |
| *5HT-1B^ΔIII-V^ /5HT-1B^ΔIII-V^* | 55846 | 30% |

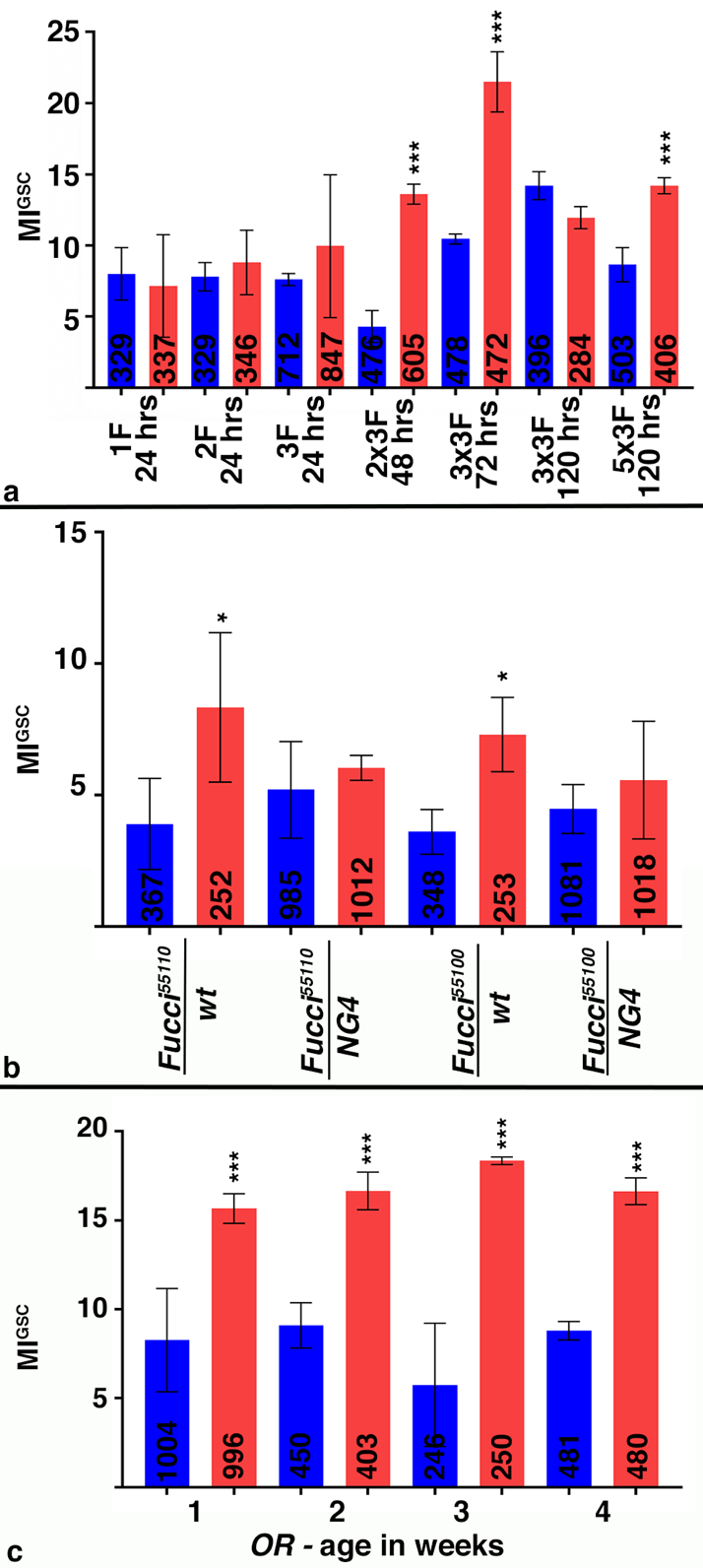

**Figure S1. Validation of mating conditions for the increase in MI^GSC^.**

A-C) Bar graphs showing MI^GSC^. Blue: non-mated condition, red: mated condition, ***: P-value < 0.001, *: P-value < 0.05, numbers of GSCs as indicated, genotypes as indicated.

A) MI^GSC^ after different mating conditions, as indicated. F: female virgins, hrs: hours.

B) MI^GSC^ from experimental (*Fucci/NG4*) and control (*Fucci/wt*) Fucci-lines.

C) MI^GSC^ of non-mated and mated *OR* males at one, two, three, and four weeks of age.

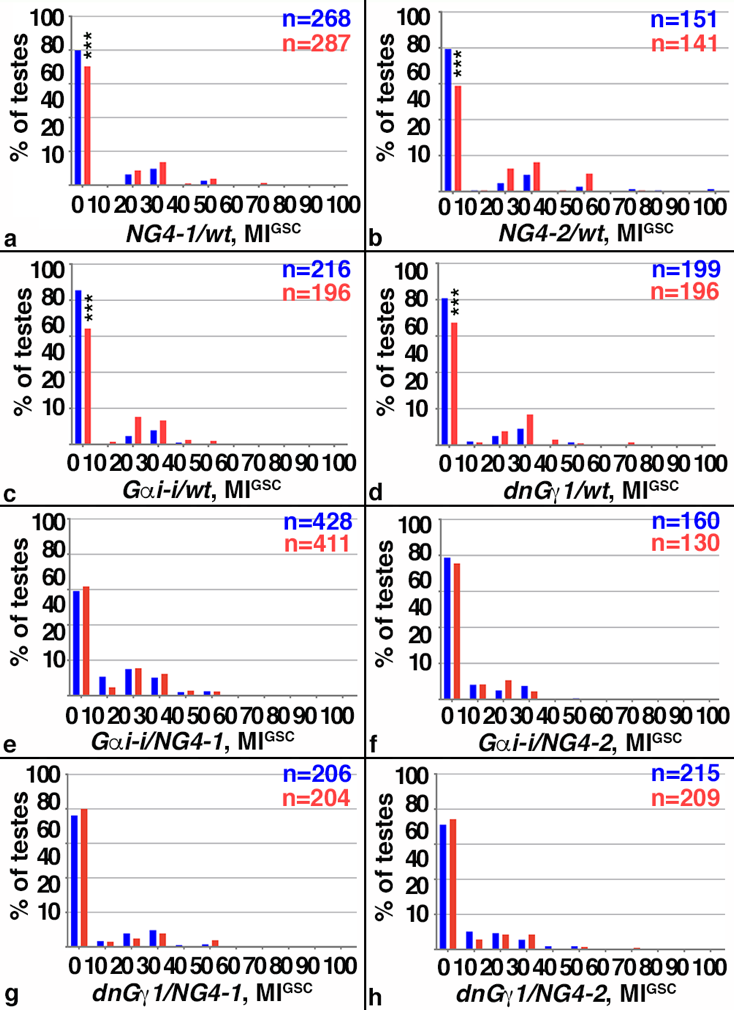

**Figure S2. Modulated G-protein did not significantly change the distribution of MIGSC across the population of testes.**

A-H) FDGs showing median of bin of MI^GSC^ across populations of males on the X-axis (bin width=10) and the percentage of testes with each MI^GSC^ on the Y-axis. Blue: non-mated condition, red: mated condition, n: number of testes examined, genotypes as indicated, ***: P-value < 0.001.

A-D) Control males

E-H) Males expressing *G_α_i-i* or dnG_γ_1 in the germline.

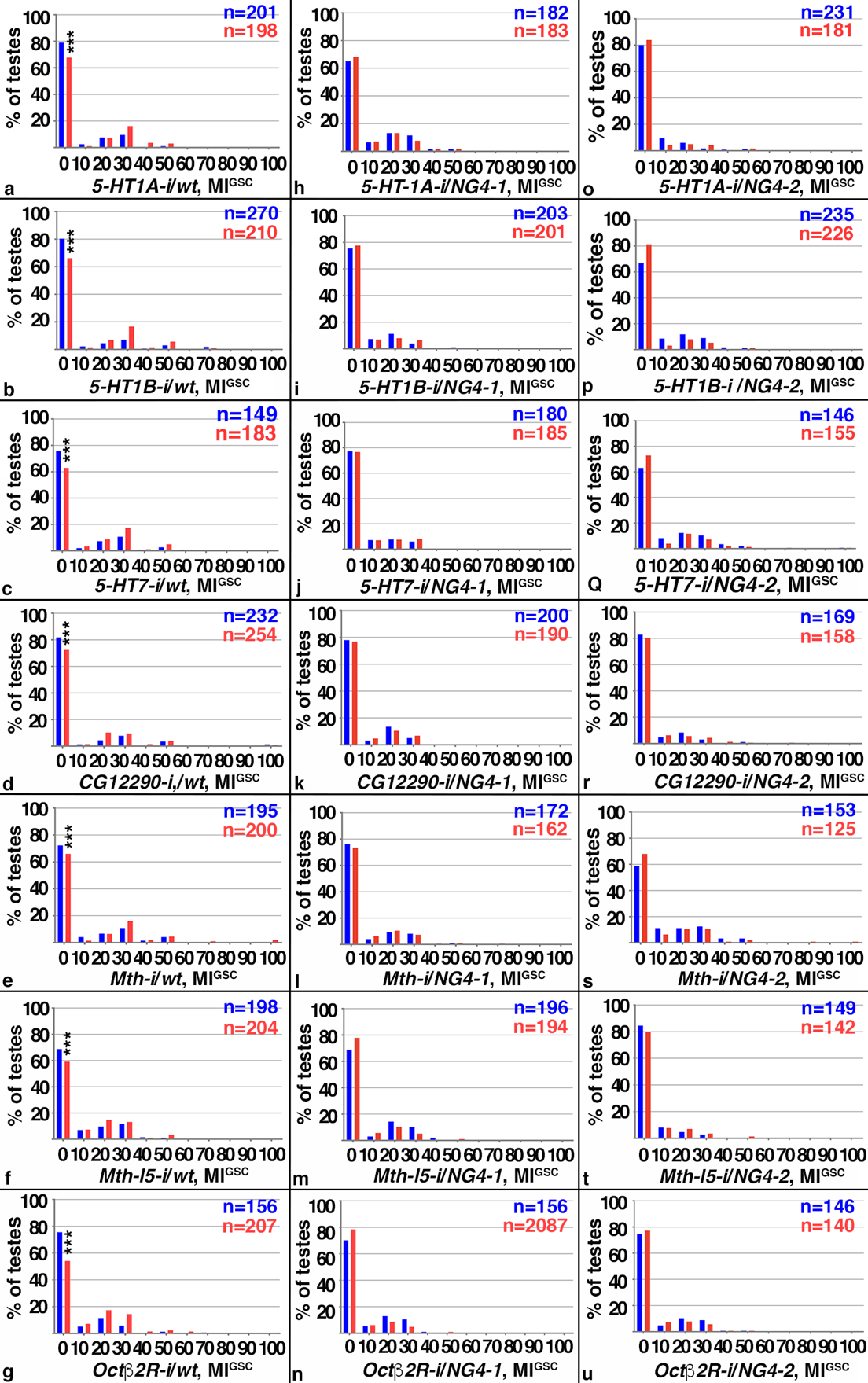

**Figure S3. No change in the distribution of MI^GSC^ in response to mating was seen upon expressing RNA-*i* against seven of the GPCRs.**

A-U) FDGs showing median of bin of MI^GSC^ across populations of males on the X-axis (bin width=10) and the percentage of testes with each MI^GSC^ on the Y-axis. Blue: non-mated condition, red: mated condition, n; number of testes examined, genotypes as indicated, ***: P-value < 0.001.

A-G) Control males.

H-U) Males expressing RNA-*i* against seven GPCRs in the in the germline via (H-N) *NG4-1* or (O-U) *NG4-2*.
